# Fibroblast growth factor-2 prevents synaptic pathology in minimal hepatic encephalopathy via NRG1/ErbB4 signaling

**DOI:** 10.1101/869503

**Authors:** Jian Wang, Weishan Zhuge, Xiaoai Lu, Ruimin You, Leping Liu, He Yu, Yiru Ye, Xuebao Wang, Qichuan Zhuge, Saidan Ding

## Abstract

**Background:** Minimal hepatic encephalopathy (MHE) is implicated in the impairment of memory function. Fibroblast growth factor-2 (FGF2) is involved in modulating synaptic and neuronal formation.

**Methods:** The aim of this study is to examined the impacts of FGF2 on MHE pathology. Our study addressed whether FGF2 could trigger neuregulin-1 (NRG1) release to ameliorate synaptic impairment in MHE rats and in primary cultured neurons.

**Results:** The results showed the decreased FGF2 expression in MHE brains. After treatment with FGF2, secreted neuregulin-1 (NRG1) and ErbB4 were increased, and the interaction of the 2 proteins was enhanced. Additionally, treatment with FGF2 or NRG1 induced synaptic formation, with increase in the activity of synapse and the density of dendritic spine, through Sirt1. NRG1 signaling was prevented by administration of FGF2, which acts through the FGFR1 in MHE rats. Finally, intracerebroventricular injection with FGF2 or NRG1 mitigated the impairment of synaptogenesis.

**Conclusion:** The data suggest that FGF2 may be a promising latent therapeutic reagent for MHE pathogenesis.

## Introduction

Fibroblast growth factor-2 (FGF2) is most abundantly expressed in the central nervous system (CNS) by adulthood (*1*). Studies on rodents have found that released FGF2 primarily binds to FGF receptor-1 (FGFR1) located on the cell surface, though can interact with all 4 FGF receptors (*2*). FGF2 is implicated in regulating many functions of CNS, such as, enhances neuronal survival and neurogenesis, facilitates dendritic growth (*3, 4*) and regenerative plasticity (*5, 6*), and promotes synaptic formation (*7*). Minimal hepatic encephalopathy (MHE) mainly is characterized by disruption in memory function detectable by electrophysiological tests (*8*). However, the role of FGF2 in MHE pathology is unknown. We recently proposed that FGF2 might also be involved in modulating synaptogenesis underlying cognitive function in MHE rats.

Neuregulin-1 (NRG1), is a member of an active epithelial growth factor (EGF) family, identified as a transmembrane protein. NRG1 and its ErbB receptor tyrosine kinases, are expressed in the CNS, where NRG1 binds to and activates ErbB. They exert an essential role in regulating the synaptic plasticity, regeneration of the nervous system (*9–11*) and homeostasis of brain activity (*12*). An abundant evidence suggests that NRG1 has been implicated in preventing brain injury following stroke (*13, 14*) and protecting neurons in Parkinson’s disease (*15*) and Alzheimer’s disease (AD) (*16*). Therefore, FGF2 may regulate the NRG1/ErbB4 pathway involved in synaptic formation in MHE.

Prior to the present study, the expression of FGF2 and its effect on cognitive function—which is associated with NRG1 secretion in MHE rats—had not been investigated. We hypothesized that reducing FGF2 levels in MHE rats would prevent cognitive decline by impairing the activation of NRG1/ErbB4 signaling and dowregulation of Sirt1.

## Materials and methods

### MHE models and treatment

Sprague-Dawley rats (220-250g) were purchased from the experimental animal center of the Chinese academy of sciences in Shanghai. The normal values of water-finding task (WFT) and Y-maze (YM) of all animals were achieved before experimenting. To induce liver cirrhosis, animals (n=30) were injected (i.p.) with thioacetamid (TAA) (200 mg/kg in normal saline, Sigma-aldrich) twice per week for 8 weeks. TAA-treated rats were included to the HE group, if they exhibited symptoms of delayed motor activity, lethargy, and subsequent coma (*17*). TAA-treated rats with no HE symptoms (n=22) were conducted to the WFT and YM to confirm MHE condition with the requirements for values of WFT higher than mean ±1.96·SD or values of YM lower than mean ±1.96·SD.

Normal and MHE rats were implanted with cannulae in the right lateral ventricles using stereotaxis instrument. MHE rats were intracerebroventricularly microinjected with FGF2 (0.6 μg/10 μl, 1.2 μg/10 μl) or NRG1 (5 μmol/5 μl, 10 μmol/5 μl) through a guide cannula for 3 times one week, or with 0.5 μg of FGF2, NRG1 plasmid or control plasmid pCMV-Tag2A for 24 h (Santa Cruz, CA, USA) (*18*). After injection, YM and WFT tests were conducted.

### Behavioral tests

For YM, rats were individually put at the end of an arm in a three arm apparatus to explore the maze freely for 8 min. Spontaneous alternation percentage (SA%), defined as a ratio of the arm choices to total choices, were measured (*19*).

For WFT, Rats were individually put at the near-right corner of the WFT apparatus and allowed to find and drink the water in the alcove within 3min. Entry latency (EL, the elapsed times for entry into the alcove), contacting latency (CL, the elapsed times for the first touching/sniffing/licking of the water tube) and drinking latency (DL, the elapsed times for the initiation of drinking from the water tube) were measured (*20, 21*).

### Cell culture and treatments

Primary hippocampal rats neurons (PHNs) and Primary cortical rats neurons (PCNs) were obtained from respective hippocampus or cerebral cortex of 1-day-old Sprague-Dawley rat pups which were digested with trypsin and DNase, then plated in poly-L-lysine-precoated six-well plates with a density of 2×106 cells/well in Neurobasal® Medium (1X) supplemented with 0.5 mM GlutaMAX™-I, B-27®(*22*). Then PHNs or PCNs were treated with FGF2 or NRG1 (1, 5, 20 ng/ml) for 24 h in the presence or absence of 10 μg/ml polyclonal anti-NRG1 antibody (Origene, Rockville, MD), 50 μmol/L ErbB4 inhibitor AG1478 (Origene, Rockville, MD), or 10 μmol/L Sirt1 inhibitor sirtinol (Origene, Rockville, MD) for 24h, or treated with FGF2 (20 ng/ml) for 24 h after transfection with Silencer Negative Control #1 siRNA (scrambled siRNA), NRG1 or ErbB4 siRNA (0.25 μg, Santa Cruz, CA, USA).

### RT-PCR and real-time quantitative PCR (qPCR)

cDNAs templates were synthesized from extracted total RNA using omniscript reverse transcriptase (Quiagen) and used to perform PCR amplification under the indicated conditions using Taq DNA polymerase (Sigma-Aldrich).

Real-time PCR results were normalized to GAPDH mRNA. The primers (Invitrogen) were as follows: NRG1, 5’AATGGACAGCAACACAAG3′ (Forward) and 5’TTAGCGATTACACTAGACAG3′ (Reverse); FGF2, 5’GAAGAGCGACCCTCACATCAAG3′ (Forward) and 5’CTGCCCAGTTCGTTTCAGTG3′ (Reverse); GAPDH, 5′TGTCATCAACGGGAAGCCCA3′ (Forward) and 5’TTGTCATGGATGACCTTGGC3′ (Reverse).

### Measurement of FGF2 or NRG1 release

High sensitivity sandwich enzyme-linked immunosorbent assay (ELISA) kits were used to assay Extracellular FGF2 or NRG1 levels in the culture medium of primary neurons in 96-well plates. FGF2 or NRG1 levels were determined spectrophotometrically using a Thermo-Fisher Multiskan MCC plate reader according to the manufacturers’ recommendations.

### Immunoblotting (IB) analysis

The amount of protein was prepared from homogenized tissues or tells and determined using the Bradford quantification assay (Bio-Rad). Protein extracts were separated by SDS-PAGE and electroblotted to PVDF membrane (Millipore, Bedford, MA, USA). After blocking with 5% (w/v) non-fat dry milk in PBS, the membrane was probed overnight at 4°C with primary antibodies (ErbB4, pErbB4, NRG1, FGF2, Sirt1, syntaxin, Homer, or β-actin) (Abcam, Cambridge, UK), and incubated with horseradish peroxidase-conjugated secondary antibodies (Pierce) for 1 h at room temperature. Blots were developed by ECL reagent (Amersham, Arlington Heights, IL, USA), and recorded on Kodak Biomax film.

For coimmunoprecipitations, lysates of tissues or tells were incubated with antibodies overnight (4°C) and subsequently with protein G-agarose beads (Millipore) for 5h (4 °C). Beads were washed with lysis buffer, the eluent was separated by SDS-PAGE and electroblotted to PVDF membrane to probe proteins using primary and secondary antibodies.

### Functional labeling of presynaptic boutons with FM4-64

5 mg/mL FM4-64 (Invitrogen) and 50 mM KCl in Hanks’ balanced salt solution were used to incubate primary neurons for 1 min at 4°C. After reaction, free FM4-64 was removed by washing with Hanks’ balanced salt solution.

### Double-labeled fluorescent staining

Brain sections or glass coverslips were fixed with 4 % paraformaldehyde for 30 min, rupture of cell membrane was performed using 0.1% Triton X-100 for 10 min at room temperature, PBS containing 5% normal goat serum for 1 h at room temperature was used for blocking, and sections or coverslips were incubated overnight at 4 °C with the following primary antibodies: pErbB4, NRG, FGF2, Sirt1, syntaxin, Homer, and MAP2 (Abcam). Alexa Fluor 488 (green)/Alexa Fluor 594 (red) conjugated secondary antibody (Abcam, Cambridge, MA) were then used to detect primary antibodies for 1h.

For dendritic spine analysis, the primary neurons coverslips were incubated with the primary antibodies: microtubule-associated protein 2B (MAP2B; 1:200; BD Transduction Laboratories, San Jose, CA, USA) and vesicular glutamate transporter 1 (vGlut1; 1:100; Neuromab, Davis, CA, USA) overnight at 4°C. Alexa Fluor 488 (green)/Alexa Fluor 594 (red) conjugated secondary antibody (Abcam, Cambridge, MA) were then used to detect primary antibodies for 1h. At least 10 cultured primary neurons per coverslip were used for quantitative analysis.

### Statistical analysis

All of the data were indicated as mean ± SD. Data comparisons were determined using one-way analysis of variance (ANOVA). Dunnett’s post hoc multiple comparison test was performed when significant differences were achieved by the ANOVA model. Then P values were adjusted by Bonferroni correction. The level of significance was determined for P < 0.05 or P < 0.01. All analyses were performed with SPSS 18.0 (PASW Statistics 18.0).

## Results

### FGF2 was reduced in the brain of the MHE rat model

FGF2 has been reported to be implicated in synaptogenesis and neuroprotection (*23*). Based on this finding, we examined the expression of FGF2 in MHE rats. Through IB analysis, the decrease in the expression of FGF2 levels were observed in the hippocampus and cortex of MHE rats (Fig. 1a and b). By IF staining, we also confirmed the reduced expression of FGF2 in cortex of MHE rats (Fig. 1c). RT-PCR (Fig. 1d) and qPCR (Fig. 1e) also showed the reduction of FGF2 transcription levels in the hippocampus and cortex of MHE rats, and ELISA assay showed the decrease in FGF2 levels in the hippocampus and cortex of MHE rats (Fig. 1f).

**Figure 1.**
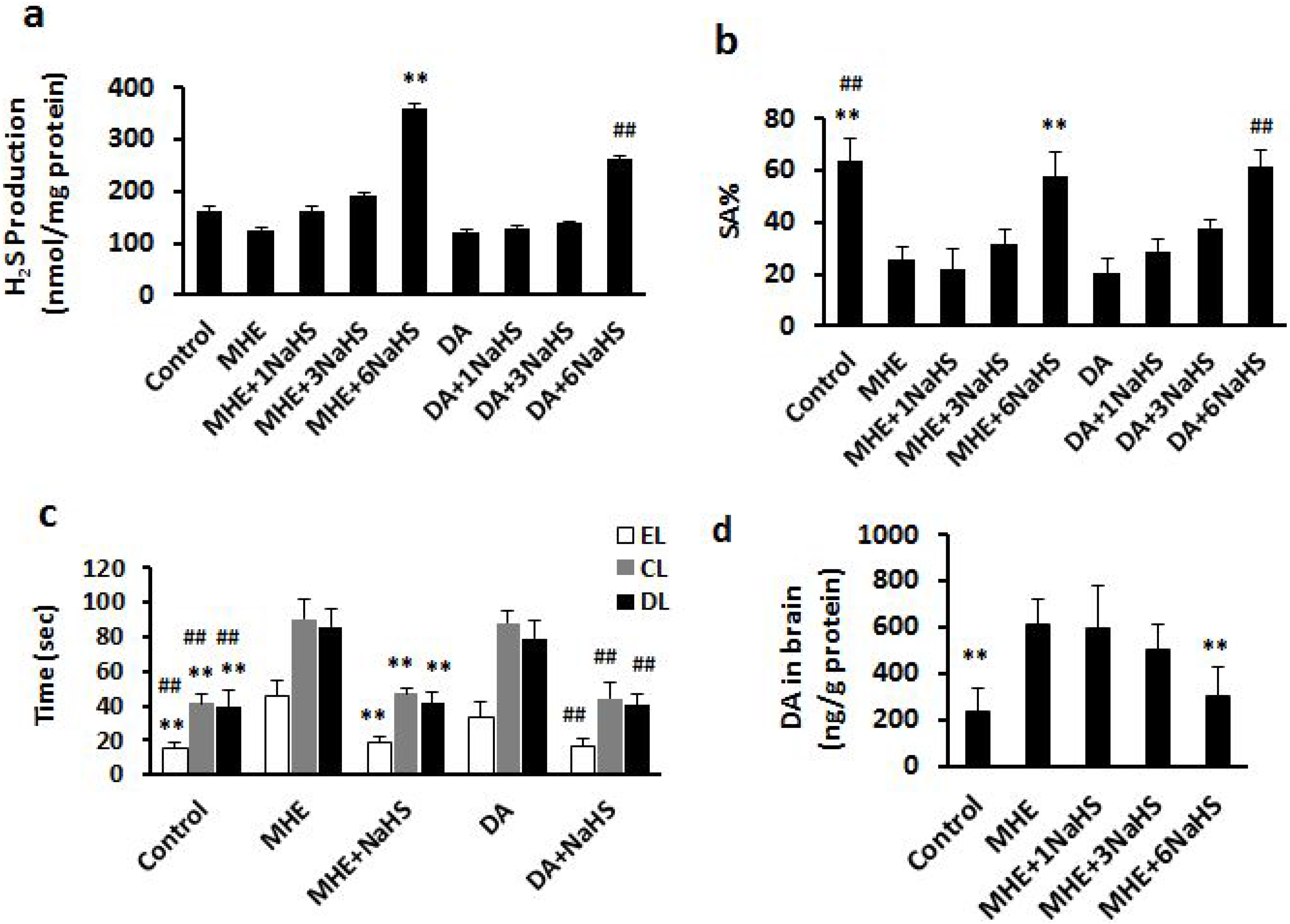
FGF2 expression was reduced in the hippocampus and cortex in MHE rats. (a, b) IB analysis of hippocampal and cortical lysates from MHE rats using (a) antibodies against FGF2 and GAPDH and (b) subsequent densitometry. (c) Immunostaining of free-floating cortical sections from MHE rats using antibodies against FGF2 (red) and MAP2 (green). (d, e) Analysis of FGF2 mRNAs of hippocampal and cortical lysates from MHE rats by (d) RT-PCR and (e) qPCR. (f) ELISA assay for FGF2 level of hippocampal and cortical lysate from MHE rats. Data are shown as mean ± SD. \**P* < 0.05, \*\**P* < 0.01 vs. MHE model group. Scale bar, 25 μm. MRGD, merged image

### FGF2 elevated NRG1 release from neurons

NRG1 was previously found to be involved in synaptic plasticity (*24, 25*). We hypothesized that FGF2 was implicated in the activation of NRG1 to regulate synaptic plasticity, and we set out to test the effect of FGF2 on NRG1 expression. First, we tested whether the FGF2 signaling extended to activate NRG1-ErbB4 signaling by neurons. The results of IB analysis, shown in Fig. 2a and b, indicated that maximal increases in NRG1 expression were achieved by 20 ng/ml FGF2, whereas a low dose of FGF2 did not alter the protein expression in PHNs, likewise, PCNs treated with FGF2 showed significant increase in NRG1 levels by 20 ng/ml (Fig. 2a and b). FGF2 treatment in PHNs significantly increased NRG1 levels by 24 hours; the same result was found in PCNs (Fig. 2c and d). IF staining confirmed the increased expression of NRG1 in PHNs in response to FGF2 (Fig. 2e). Using ELISA, the results showed that FGF2 treatment elevated the level of NRG1 release with a concentration- (Fig. 2f) and time-dependence (Fig. 2g) in PHNs and PCNs. Through RT-PCR, we found that FGF2 triggered the transcriptional process of NRG1 expression dose-dependently (Fig. 2h) and time-dependently (Fig. 2i) in PHNs and PCNs. we then examine whether FGF2-treated conditions altered the levels of other NRGs, NRG2 or NRG3. Unlike NRG1, we found that NRG2 or NRG3 levels were not changed by any concentration of FGF2 in PHNs (Fig. 2j and k).

**Figure 2.**
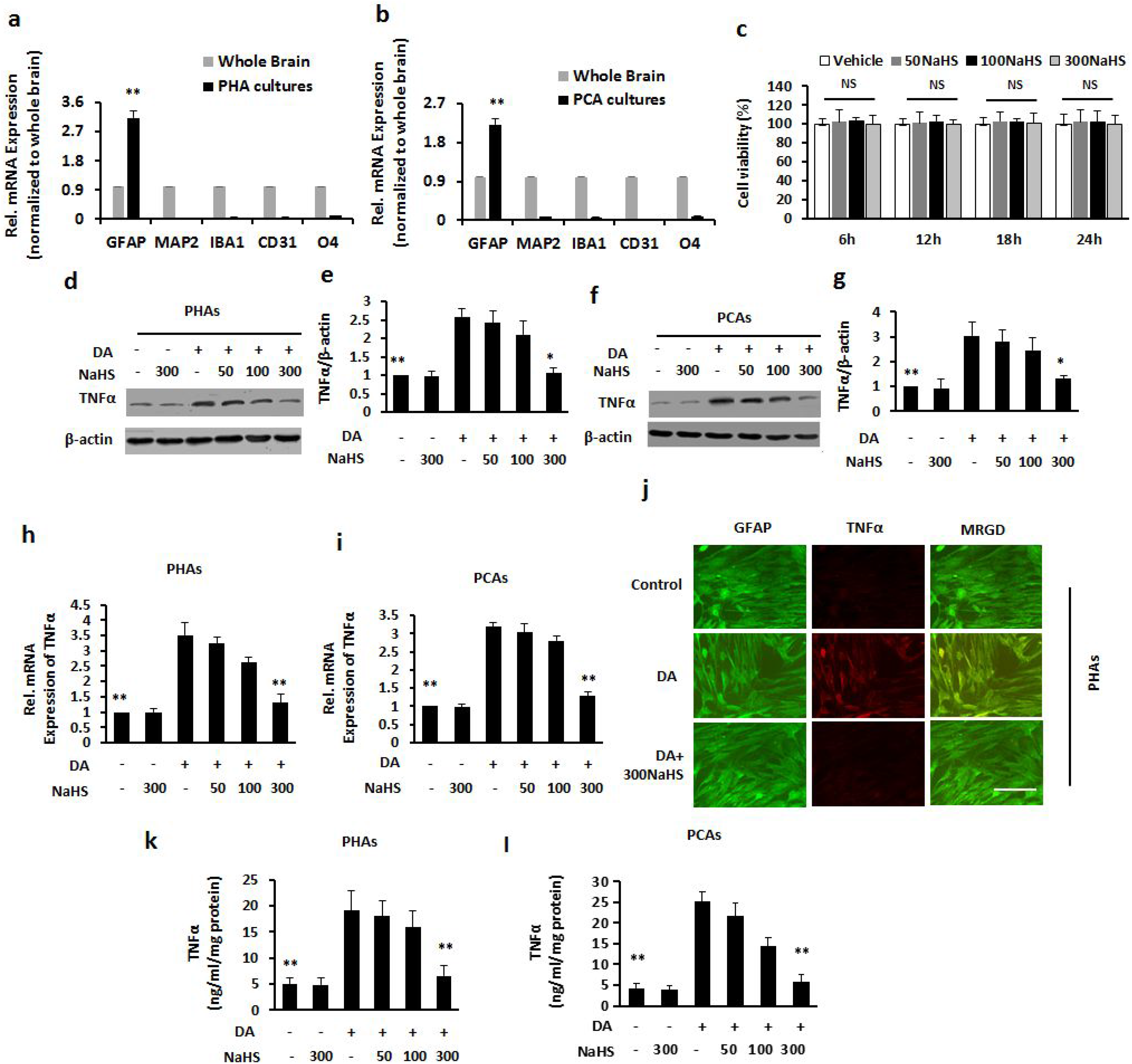
FGF2 stimulated the production and release of NRG1 *in vitro*. (a) 1B analysis of lysate from PHNs or PCNs stimulated with various concentrations of FGF2 using anti-NRG1 and anti-actin antibodies and (b) subsequent densitometry. (c) IB analysis of lysate from PHNs or PCNs stimulated with FGF2 for various time point using anti-NRG1 and anti-actin antibodies and (d) subsequent densitometry. (e) Immunostaining of PHNs stimulated with FGF2 using antibodies against NRG1 (red) and MAP2 (green). (f) RT-PCR analysis of NRG1 mRNAs of PHNs treated with different doses of FGF2. (g) RT-PCR analysis of NRG1 mRNAs of PHNs treated with 20 ng/ml FGF2 for various time points. (h) ELISA assay for RGI expression of supernatants from PHNs stimulated with various concentrations of FGF2. (i) ELISA assay for NRG1 level of supernatants from PHNs stimulated with 20 ng/ml NRG1 for various time points. (j) IB analysis of lysate from PHNs stimulated with various concentrations of FGF2 using anti-NRG2/NRG3 and anti-actin antibodies and (k) subsequent densitometry. Data are shown as mean ± SD. \**P* < 0.05, \*\**P* < 0.01 vs. vehicle-treated group. NS, not significant. Scale bar, 25 μm. MRGD, merged image.

Next, we determined the impact of FGF2 on the ErbB4 expression. Interestingly, while 20 ng/ml FGF2 significantly stimulated phosphorylation of ErbB4 in PHNs and PCNs, other concentrations did not change phosphorylation of ErbB4, according to the IB analysis (Fig. 3a and b). The phosphorylation of ErbB4 was increased by 72 h in PHNs or PCNs when neurons were treated with 20 ng/ml FGF2 (Fig. 3c and d). As shown in Fig. 3e, the IF analysis confirmed that ErbB4 was strongly expressed in PHNs with a 20 ng/ml dose of FGF2 treatment. We examined the NRG1–ErbB4 interaction in neurons using coimmunoprecipitation. As shown in Fig. 3f, NRG1 was immunoprecipitated, and FGF2 treatment obviously increased the content of NRG1 and ErbB4 coimmunoprecipitated with NRG1 in PHNs in a dose-dependent fashion. ErbB4 was also immunoprecipitated, and IB analysis showed that FGF2 induced the increase in ErbB4 and NRG1 coimmunoprecipitation with ErbB4 in neurons dose-dependently. Unlike ErbB4, ErbB2 expression was not regulated by different concentrations of FGF2 in PHNs (Fig. 3g and h).

**Figure 3.**
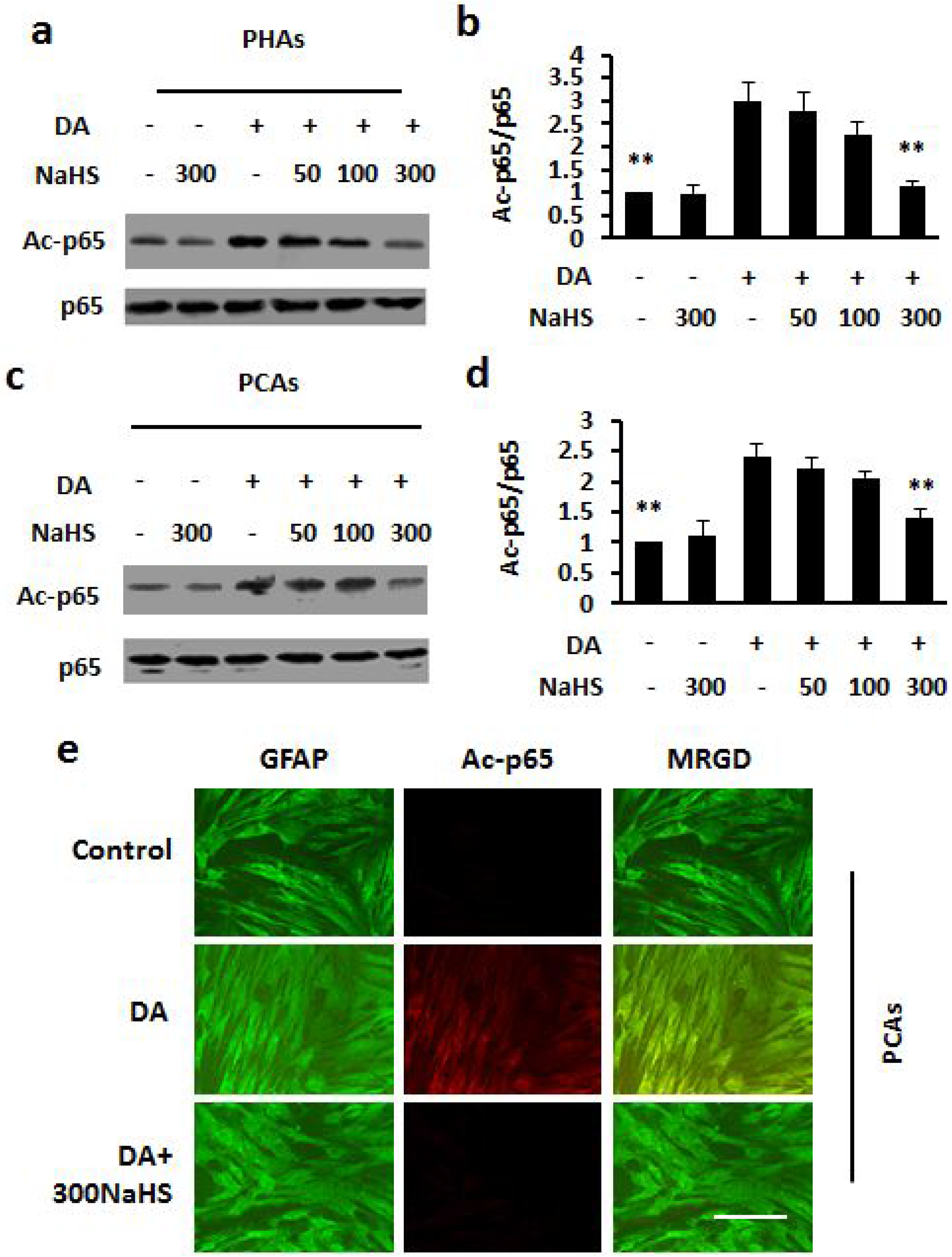
FGF2 elicited the expression of ErbB4 *in vitro*. (a) IB analysis of lysate from PHNs or PCNs stimulated with various concentrations of FGF2 using anti-pErbB4 and anti-actin antibodies and (b) subsequent densitometry. (c) IB analysis of lysate from PHNs or PCNs stimulated with FGF2 for various time points using anti-pErbB4 and anti-actin antibodies and (d) subsequent densitometry. (e) Immunostaining of PHNs stimulated with FGF2 using antibodies against pErbB4 (red) and MAP2 (green). (f) Coimmunoprecipitation analysis of lysates of cells stimulated with various concentrations of FGF2 for the association between NRG1 and ErbB4. (g) IB analysis of lysate from PHNs stimulated with various concentrations of FGF2 using anti-pErbB2 and anti-actin antibodies and (h) subsequent densitometry. Data are shown as mean ± SD. \**P* < 0.05, \*\**P* < 0.01 vs. vehicle-treated group. NS, not significant. Scale bar, 25 μm. MRGD, merged image.

### FGF2-induced synaptic formation was mediated by NRG1 signals in PHNs

Thinking that both FGF2 and NRG1 were involved in synaptic plasticity, and that FGF2 triggered release of NRG1, we investigated whether FGF2 induced synaptic activity via NRG1. As shown in Fig. 2g and h, increased syntaxin and Homer were observed in PHNs exposed to FGF2, while the addition of an anti-NRG1 antibody or AG1478 abolished these two proteins expressions (Fig. 4a and b). Immunostaining confirmed that the increase in Homer level was abated by addition of an anti-NRG1 antibody or AG1478 in PHNs (Fig. 4c). Using FM4-64 dye to probe activity-dependent synaptic vesicle recycling revealed that synaptic activity in PHNs with FGF2 treatment was markedly increased, but decreased with addition of the anti-NRG1 antibody or AG1478 (Fig. 4d and e).

**Figure 4.**
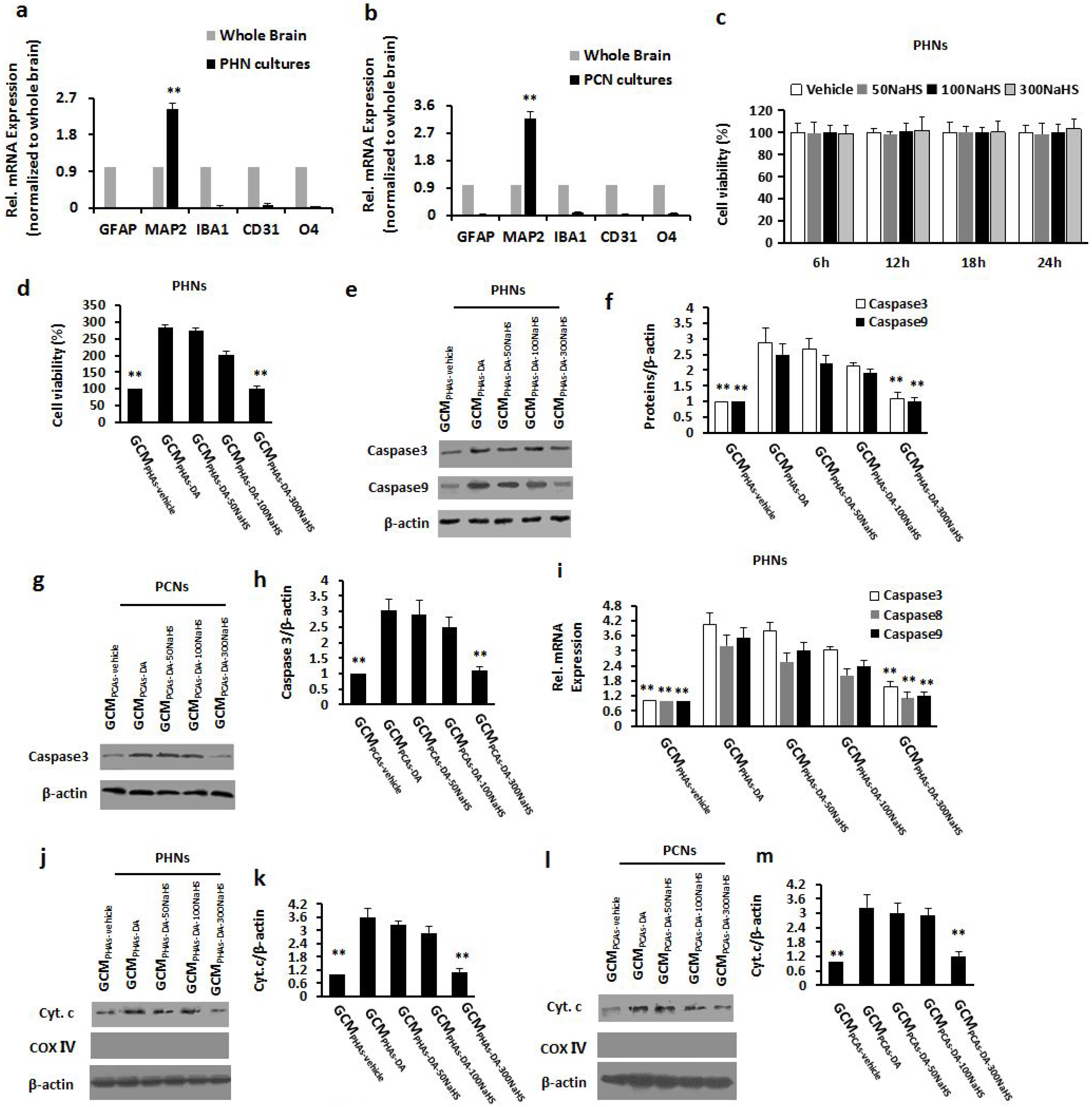
FGF2 activated NRG1/ErbB4 signaling to promote synaptic formation in primary neurons. (a) IB analysis of PHNs treated with 20 ng/ml FGF2 alone or after preincubation with anti-NRG1 antibody or AG1478 using antibodies against syntaxin/Homer and (b) subsequent densitometry. (c) Immunostaining of PHNs stimulated with FGF2 (20 ng/ml) in the presence of anti-NRG1 antibody or AG1478 using antibodies against syntaxin (red) and MAP2 (green). (d, e) Representative image of FM4-64 staining of functional presynaptic terminals in PHNs treated with 20 ng/ml FGF2 alone or after preincubation with anti-NRG1 antibody or AG1478. The (e) panel indicates that quantitative analysis of changes showed on average in FM4-64 puncta intensity. (f) IB analysis of PHNs transfected with NRG1 siRNA using anti-NRG1 and anti-actin antibodies and (g) subsequent densitometry. (h) IB analysis of PHNs transfected with ErbB4 siRNA using anti-ErbB4 and anti-actin antibodies and (i) subsequent densitometry. (j) IB analysis of PHNs transfected with NRG1 or ErbB4 siRNA in the presence or absence of FGF2 (20 ng/ml) using anti-syntaxin/Homer and anti-actin antibodies and (k) subsequent densitometry. (l, m) Immunostaining of PHNs transfected with NRG1 or ErbB4 siRNA in the presence of FGF2 against MAP2B (red) and vGluT1 (green). Red signals indicate MAP2B for microtubule staining and green signals indicate vGluT1 for detecting excitatory synapses. Synaptic density was analyzed by counting green signals (vGluT1-positive dendritic spines) using ImageJ, and expressed per 1 μm of apical dendrite. Scale bar, 2 μm. Data are shown as mean ± SD. *P < 0.05, **P < 0.01 vs. vehicle-treated group; #*P* < 0.05, ##*P* < 0.01 vs. FGF2-treated group. NS, not significant. Scr, scrambled. Scale bar, 25 μm. MRGD, merged image.

Simultaneously, PHNs were transfected with NRG1 siRNA or ErbB4 siRNA plasmids. the results of transfection efficiency showed NRG1 protein level was significantly decreased in NRG1 siRNA-transfected PHNs by IB analysis (Fig. 4f and g), and ErbB4 protein levels were also decreased in ErbB4 siRNA-transfected PHNs (Fig. 4h and i). Furthermore, we found that the FGF2-induced elevation of syntaxin and Homer was reversed by NRG1 siRNA or ErbB4 siRNA transfection in PHNs (Fig. 4j and k). The elevated Homer from high-dose FGF2 treatment was reversed by NRG1 siRNA or ErbB4 siRNA transfection in PHNs. We further addressed the role of FGF2 in the modulation of dendritic spine density using anti-vGluT1 and anti-MAP2 antibodies. Double IF staining revealed that vGluT1-positive signals were significantly elevated in PHNs exposed to NRG1 (Fig. 4l and m). These results confirmed that FGF2 induced NRG1 release and the activation of NRG1/ErbB4, leading to synaptic formation.

### Sirt1 was involved in FGF2-mediated synaptic formation

Sirt1 has been reported to be involved in cognitive function, and synaptogenesis (*26*). We tested whether FGF2 and NRG1 elicited synaptic formation via Sirt1. FGF2 significantly increased thecontent of Sirt1 dose-dependently in PHNs (Fig. 5a and b). PCNs showed the same results in the protein in response to FGF2 (Fig. 5a and b). Sirt1 levels were elevated by 48 hours in PHNs after FGF2 treatment (Fig. 5c and d). Immunostaining confirmed the elevation of Sirt1 expression in PHNs with a high dose of FGF2 (Fig. 5e). Neurons exposed to NRG1 showed dose-dependent increases in the expression of Sirt1 in PHNs and PCNs (Fig. 5f and g). Sirt1 was increased by 20 ng/ml NRG1 treatment time-dependently (Fig. 5h and i). As shown in Fig. 5j and k, the addition of sirtinol in PHNs abolished the FGF2-stimulated upregulation of syntaxin and Homer. The expressions of syntaxin and Homer were increased by NRG1, which was reversed by sirtinol (Fig. 5l and m). These results confirmed that FGF2 and NRG1 induced Sirt1 expression, leading to synaptogenesis.

**Figure 5.**
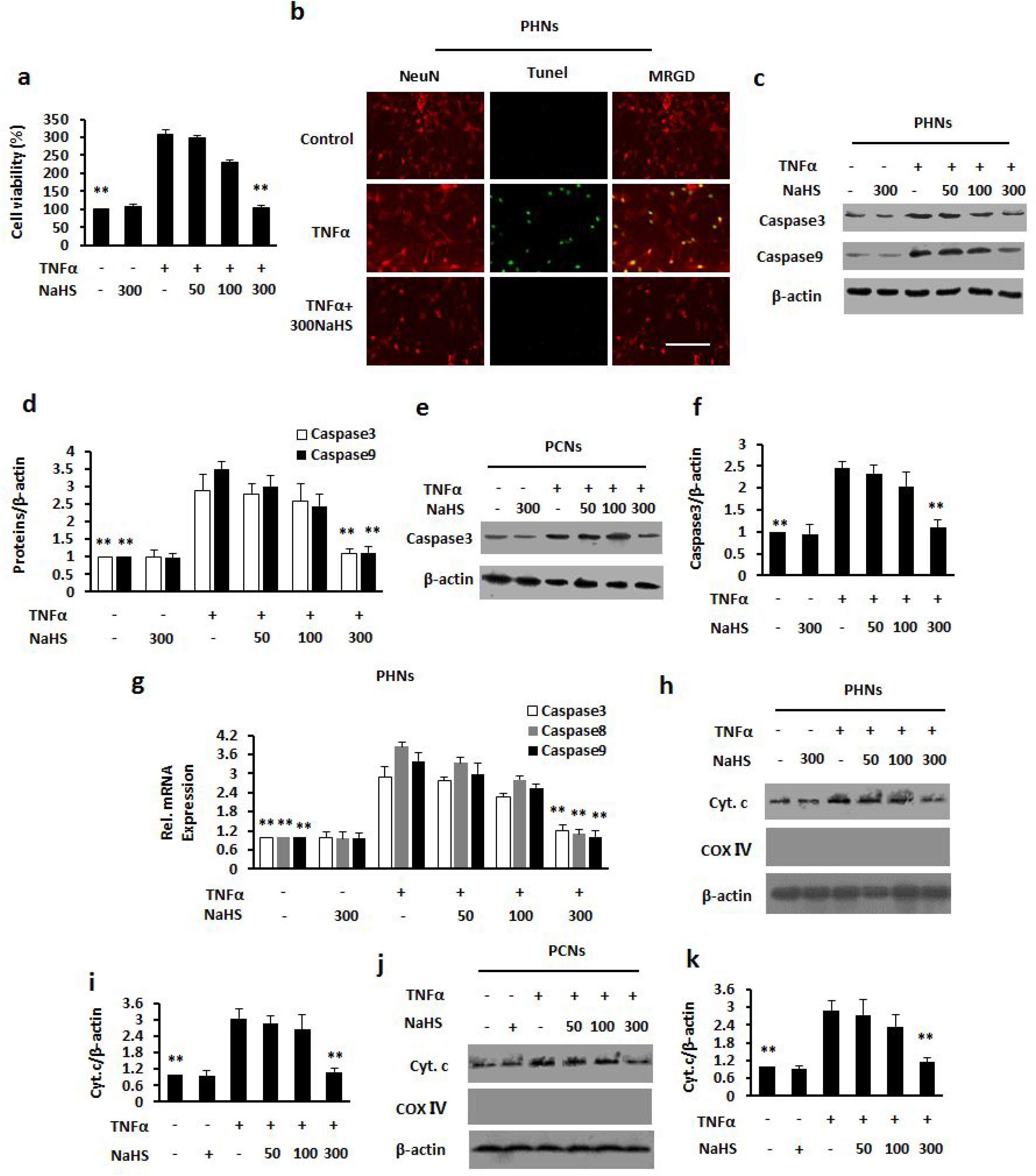
FGF2 or NRG1 upregulated Sirt1 to facilitate synaptic formation in primary neurons. (a) IB analysis of PHNs or PCNs stimulated with various concentrations of FGF2 using anti-Sirt1 and anti-actin antibodies and (b) subsequent densitometry. (c) IB analysis of PHNs treated with 20 ng/ml FGF2 for various time points using antibodies against Sirt1 and actin and (d) subsequent densitometry. (e) Immunostaining of PHNs stimulated with FGF2 (20 ng/ml) using antibodies against Sirt1 (red) and MAP2 (green). (f) IB analysis of PHNs or PCNs stimulated with various concentrations of NRG1 using anti-Sirt1 and anti-actin antibodies and (g) subsequent densitometry. (h) IB analysis of PHNs treated with 20 ng/ml NRG1 for various time points using antibodies against Sirt1 and actin and (i) subsequent densitometry. (j) IB analysis of PHNs treated with 20 ng/ml FGF2 alone or after preincubation with sirtinol using antibodies against syntaxin/Homer and (k) subsequent densitometry. (l) IB analysis of PHNs treated with 20 ng/ml NRG1 alone or after preincubation with sirtinol using antibodies against syntaxin/Homer and (m) subsequent densitometry. Data are shown as mean ± SD. \**P* < 0.05, \*\**P* < 0.01 vs. vehicle-treated group; #*P* < 0.05, ##*P* < 0.01 vs. FGF2-treated group. NS, not significant. Scr, scrambled. Scale bar, 25 μm. MRGD, merged image.

### NRG1 triggered FGF2 release from neurons

We then tested whether NRG1 in turn affected FGF2 release. IB analysis of cell lysates showed that NRG1 significantly gradually increased FGF2 in PHNs with a dose-dependent fashion, and FGF2 content in PCNs was elevated with a dosedependence in response to NRG1 (Fig. 6a and b). PHNs in response to NRG1 exhibited increases in FGF2 expression time-dependently (Fig. 6c and d). By immunostaining, NRG1 increased cell-bound FGF2 (Fig. 6e) expression in PHNs. The results of qPCR showed that FGF2 mRNA expression were increased by NRG1 dose-dependently in PHNs and PCNs (Fig. 6f). NRG1 treatment revealed steady increases in FGF2 mRNA with time-dependent fashion in PHNs. Using ELISA assay, NRG1 (20 ng/ml) elevated the release of FGF2 dose-dependently (Fig. 6h) and time-dependently in PHNs (Fig. 6i).

**Figure 6.**
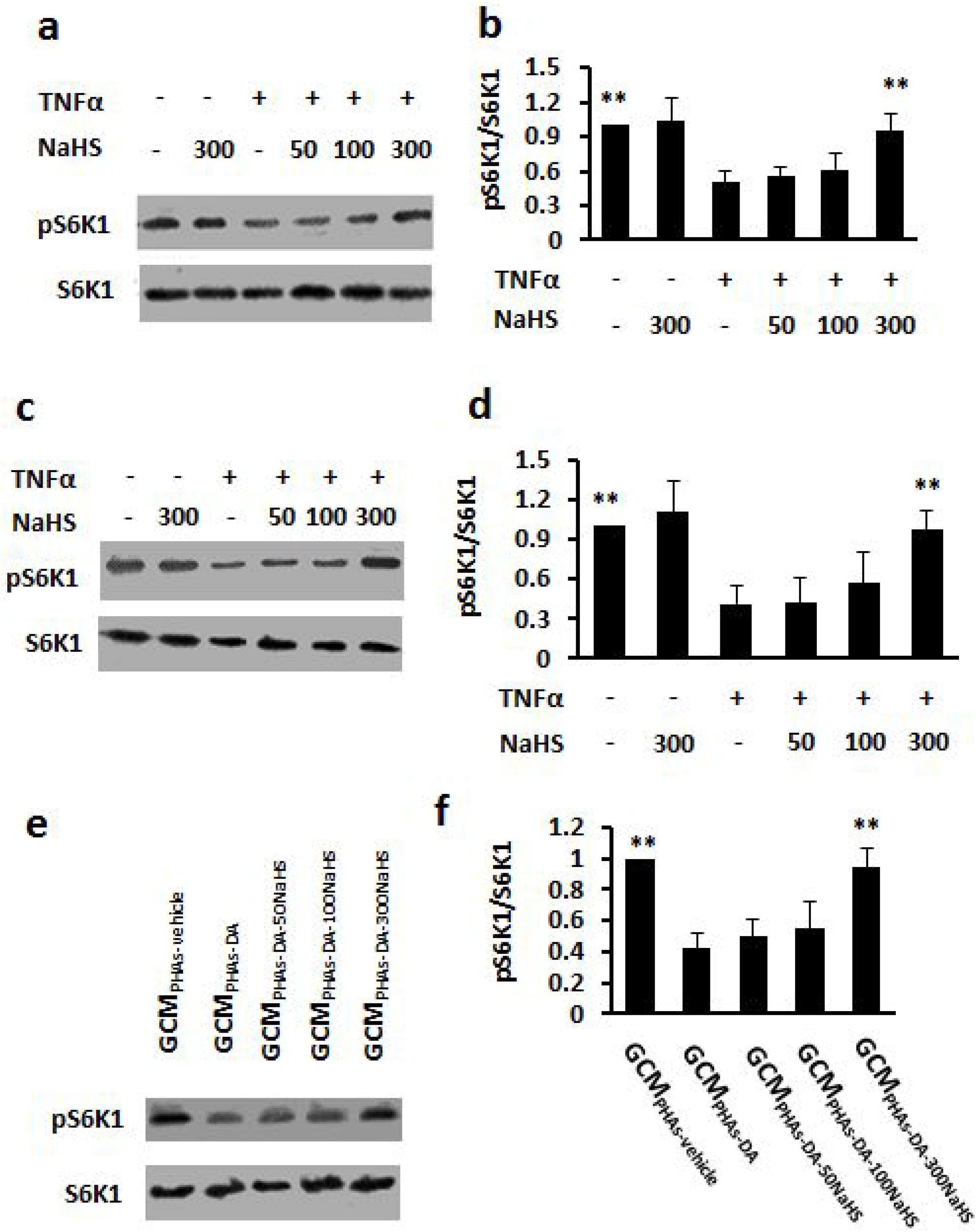
NRG1 triggered the release of FGF2 from primary neurons. (a) IB analysis of lysate from PHNs or PCNs stimulated with various concentrations of NRG1 using anti-FGF2 and anti-actin antibodies and (b) subsequent densitometry. (c) IB analysis of lysate from PHNs or PCNs stimulated with NRG1 for various time point using anti-FGF2 and anti-actin antibodies and (d) subsequent densitometry. (e) Immunostaining of PHNs stimulated with NRG1 (20 ng/ml) using antibodies against FGF2 (red) and MAP2 (green). (f) Analysis for FGF2 mRNAs of PHNs treated with different doses of NRG1 by RT-PCR. (g) Analysis for FGF2 mRNAs of PHNs treated with 20 ng/ml NRG1 by RT-PCR for various time points. (h) ELISA assay for FGF2 level of supernatants from PHNs stimulated with various concentrations of NRG1. (i) ELISA assay for FGF2 level of supernatants from PHNs stimulated with 20 ng/ml FGF2 for various time points. Data are shown as mean ± SD. \**P* < 0.05, \*\**P* < 0.01 vs. vehicle-treated group. NS, not significant. Scr, scrambled. Scale bar, 25 μm. MRGD, merged image.

### FGF2 mitigated the inactivation of NRG1/ErbB4 signaling in brain

Next, we investigated whether FGF2 had the effect on the NRG1/ErbB4 signaling *in vivo*. As determined by IB analysis in Fig. 7a and b, NRG1/ErbB4 levels were significantly reduced in the hippocampi of MHE rats, whereas, administration of high FGF2 obviously restored the expression of the 2 proteins. However, low FGF2 did not alter NRG1/ErbB4 levels in MHE rats. We also observed the decrease in NRG1/ErbB4 expression in the cortices of MHE rats, which was blocked by FGF2 administration dose-dependently (Fig. 7c and d). Immunostaining of the cortices of MHE rats showed pronounced decrease in NRG1 levels, which were diminished by administration of high FGF2 (Fig. 7e). At the same time, IB analysis showed that FGF2 protein levels were significantly increased in the hippocampi and cortices of normal rats with FGF2 gene transfection compared to normal rats and in the hippocampi and cortices of MHE rats with FGF2 gene transfection compared to MHE rats (Fig. 7f and g), indicating high transfection efficiency. Then, we found that NRG1/ErbB4 protein levels were significantly increased in the hippocampi of MHE rats, and FGF2 transfection blocked the decrease (Fig. 7h and i). The cortices of MHE rats showed decrease in NRG1/ErbB4 protein levels, which was inhibited by FGF2 transfection (Fig. 7j and k). Using IF staining, we confirmed that the strong expression of ErbB4 was induced after Wnt5a transfection (Fig. 7l). IB analysis from hippocampal lysates showed that the decrease in pFGFR1 levels in MHE rats was reversed by FGF2 administration (Fig. 7m and n). The pFGFR1 levels were reduced in the cortices of MHE rats (Fig. 7o and p), while FGF2 transfection blocked this protein. These data suggest that the administration of FGF2 activated the NRG1/ErbB4 signaling by the brain of MHE rats.

**Figure 7.**
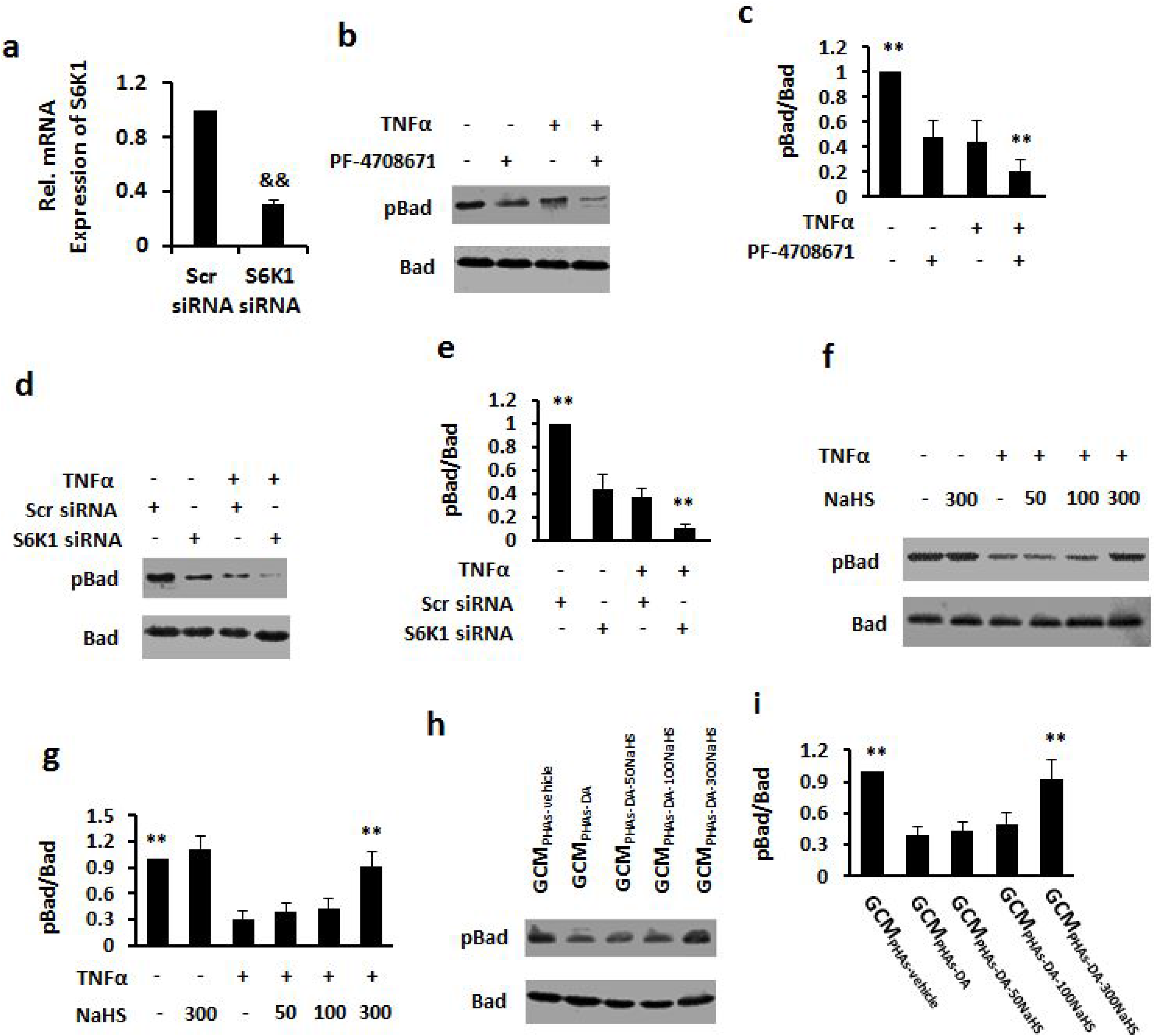
FGF2 blocks the inactivation of NRG1/ErbB4 signaling *in vivo*. (a–d) IB analysis and subsequent densitometry of hippocampal (a, b) and cortical (c, d) homogenates from MHE rats administered (ICV) with various concentrations of FGF2 using antibodies against NRG1/ErbB4 and actin. (e) Immunostaining of free-floating cortical sections from MHE rats administered (ICV) with FGF2 (1.2 μg/10 μl) using antibodies against NRG1 (red) and MAP2 (green). (f) IB analysis of hippocampal or cortical homogenates from control or MHE rats with FGF2 transfection using antibodies against FGF2, and ACTIN and (g) subsequent densitometry. (h–k) IB analysis of hippocampal (h, i) and cortical (j, k) homogenates from control or MHE rats with FGF2 transfection using antibodies against NRG1/ErbB4 and actin and subsequent densitometry. (l) Immunostaining of free-floating cortical sections from MHE rats with FGF2 transfection using antibodies against ErbB4 (red) and MAP (green). (m) IB analysis of hippocampal homogenates from MHE rats administered (ICV) with various concentrations of FGF2 using antibodies against pFGFR1 and actin and (n) subsequent densitometry. (o) IB analysis of hippocampal homogenates from MHE rats with FGF2 transfection using antibodies against pFGFR1 and actin and (p) subsequent densitometry. Data are shown as mean± SD. \**P* < 0.05, \*\**P* < 0.01 vs. control group; #*P* < 0.05, ##*P* < 0.01 vs. MHE model group. NS, not significant. Scr, scrambled. Scale bar, 25 μm. MRGD, merged image.

### FGF2 attenuated the disruption of synaptic formation *in vivo*

We assessed the effect of FGF2 on synaptic formation. As indicated by the IB analysis of the hippocampi of MHE rats showed the reduction of syntaxin and Homer expression, and FGF2 administration resulted in an increase in the 2 proteins; NRG1 administration also diminished the proteins in. Syntaxin/Homer expression (Fig. 8c and d) was significantly decreased in the cortices of MHE rats, whereas, administration of FGF2 and NRG1 reversed the expression of the 2 proteins. As shown by IF staining, MHE rats displayed lower expression of Homer, and administration of NRG1 increased the expression of this protein, while administration of FGF2 diminished the decrease in the protein (Fig. 8e).

**Figure 8.**
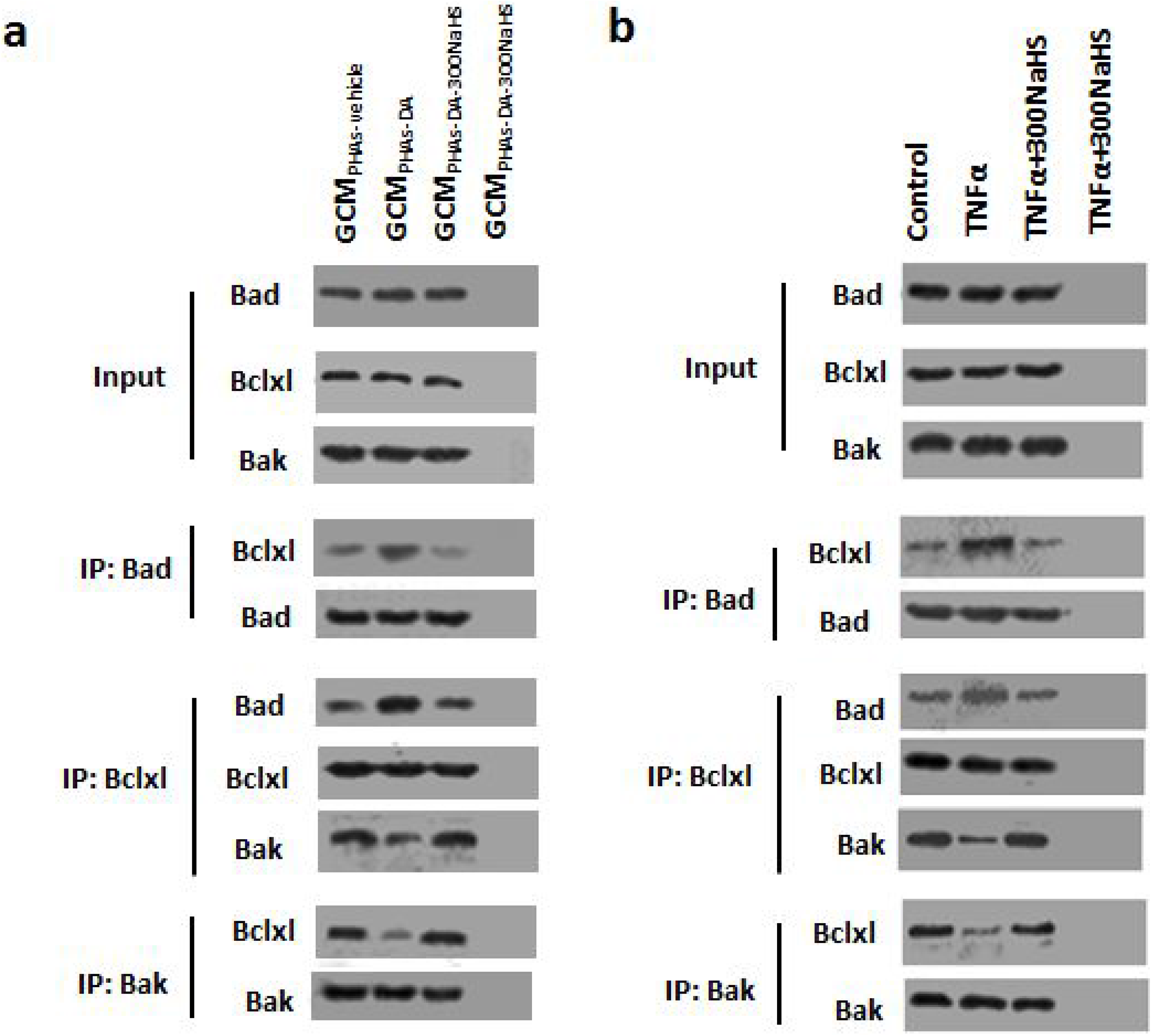
FGF2 or NRG1 attenuated the disruption of synaptic formation in MHE rats. (a–d) IB analysis and subsequent densitometry of hippocampal (a, b) and cortical (c, d) homogenates from MHE rats administered (ICV) with various concentrations of FGF2 or NRG1 using antibodies against syntaxin/Homer and actin. (e) Immunostaining of free-floating cortical sections from MHE rats administered (ICV) with FGF2 (1.2 μg/10 μl) or NRG1 (10 μmol/5 μl) using antibodies against Homer (red) and MAP2 (green). (f) IB analysis of hippocampal homogenates from control or MHE rats with NRG1 transfection using antibodies against FGF2 and actin and (g) subsequent densitometry. (h) IB analysis of hippocampal homogenates from control or MHE rats with FGF2 or NRG1 transfection using antibodies against syntaxin/Homer and actin and (i) subsequent densitometry. (j, k) Immunostaining of free-floating cortical sections from MHE rats with FGF2 or NRG1 transfection using antibodies against (j) syntaxin and (k) Homer (red) and MAP2 (green). Data are shown as mean ± SD. \**P* < 0.05, \*\**P* < 0.01 vs. control group; #*P* < 0.05, ##*P* < 0.01 vs. MHE model group. NS, not significant. Scr, scrambled. Scale bar, 25 μm. MRGD, merged image.

Similarly, IB analysis showed that NRG1 protein levels were obviously increased in the hippocampi and cortices of normal rats with NRG1 gene transfection compared to normal rats and in the hippocampi and cortices of MHE rats with NRG1 gene transfection compared to MHE rats (Fig. 8f and g). Then, we found that syntaxin/Homer expressions were decreased in the hippocampi of MHE rats, which was blocked after FGF2 transfection; NRG1 transfection also abated the decrease in the 2 proteins (Fig. 8h and i). IF analysis showed that the decrease in syntaxin expression in cortices of MHE rats was diminished by FGF2 transfection or NRG1 transfection (Fig. 8j), FGF2 transfection facilitated higher expression of Homer than that of cortice of MHE rats; NRG1 transfection had the same effect (Fig. 8k). These results suggest that FGF2 improved the production of synaptic proteins in the brain of MHE rats.

### FGF2 prevented memory impairment in MHE rats

We examined whether FGF2 or NRG1 impacted on cognitive function *in vivo*. For YM test, the SA% of MHE rats was significantly decreased, and the decrease was reversed by NRG1 or FGF2 administration (Fig. 9a). NRG1 or FGF2 transfection to MHE rats significantly increased SA% (Fig. 9b). For WFT test, EL, CL, and DL were obviously increased in MHE rats, which were recovered to the normal level by administration of NRG1 or FGF2 (Fig. 9c). NRG1 or FGF2 transfection to MHE rats markedly restored EL, CL, and DL (Fig. 9d). These results suggest that the administration of FGF2 or NRG1 reversed the cognitive deficit observed in MHE rats.

**Figure 9.**
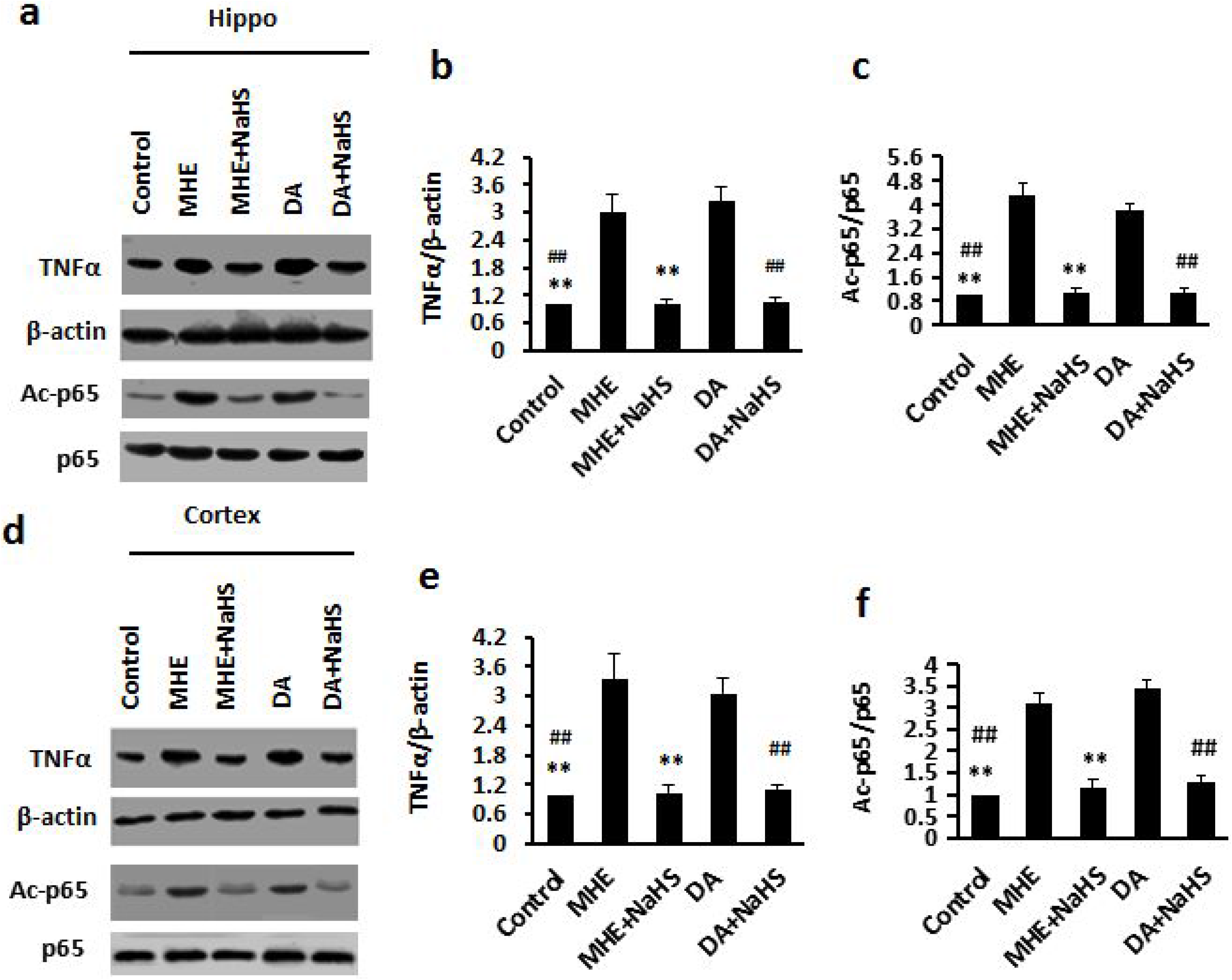
Effects of FGF2 or NRG1 on learning and memory of MHE rats. (a) Spontaneous alternation percentage (SA%) in YM of MHE rats administered with FGF2 (1.2 μg/10 μl) or NRG1 (10 μmol/5 μl). (b) SA% in YM of MHE rats with Flag-FGF2 or Flag-NRG1 transfection. (c) Results of WFT (EL, entry latency; CL, contacting latency; DL, drinking latency) of MHE rats administered with FGF2 or NRG1. (d) Results of WFT of MHE rats with Flag-FGF2 or Flag-NRG1 transfection. Data are shown as mean ± SD. \**P* < 0.05, \*\**P* < 0.01 vs. control group; #*P* < 0.05, ##*P* < 0.01 vs. MHE model group. NS, not significant. Scr, scrambled. Scale bar, 25 μm. MRGD, merged image.

## Discussion

Our present study identified NRG1/ErbB4 as a primary downstream effector of FGF2, essential for the signaling of synaptic formation. FGF2 signals mediated the activation of NRG1/ErbB4 via FGFR1, and the process was linked to the expression of Sirt1 proteins, which are sensitive to synaptic formation in neurons. We found that downregulation of FGF2 in neurons induced the inactivation of NRG1/ErbB4 signaling and downstream Sirt1, which disrupted synaptic formation, leading to the pathogenesis of MHE.

Despite the beneficial effects of FGF2 on neuron protection, synaptic plasticity, and memory function, little is known about its relationship with the pathogenesis of MHE. Our results showed that FGF2 administration immediately facilitates synaptic transmission and memory function in MHE rats. Together, our evidence supported that FGF2 played a key role in modulating synaptogenesis through the molecular cascade. our study revealed an FGF2-dependent synaptic response in neurons.FGF2 can bind to all 4 FGF receptors, located on the cell surface, but interacts with the main receptor FGFR1 (*2*). Compelling evidence showed that synaptic formation is critically facilitated by FGF2. Studies have shown that reduced FGFR1 in MHE rats was consistent with FGF2 levels. We found that decrease in FGF2 levels, but no alteration of expressions of the other FGF subfamily protein in MHE brains. The most notable finding was that the synapses were fully strengthened by administration of FGF2. The disruption of synaptic formation and the impairment of memory function were improved by FGF2 administration in MHE rats.

NRG1 is characterized by ligands of the ErbB receptor family. (*27, 28*). NRG1 possesses \an epidermal growth factor domain that is required for binding and activating ErbB3 or ErbB4 (*29*)., The adult brain widely expressed NRG1 and ErbB kinases (*30*). Furthermore, the enrichment of postsynaptic ErbB4 in particular is at excitatory synapses (*31, 32*). FGF2 protein in neurons leads to increased NRG1/ErbB4 activation. Activation of ErbB4 by binding to NRG1 affects a number of neural development, including neurotransmission, neuroplasticity (*33, 34*), or pyramidal neuron synaptic plasticity (*32, 35, 36*), and therefore impacts on behavior alteration. We wanted to identify the mechanism for the induction of MHE through the reduction of FGF2 levels. In this study, we found that FGF2 is involved in regulating the NRG1 production and secretion involved in synaptic formation, and that the concentration of NRG1 is altered in the brains of MHE rats. These findings suggest that FGF2 induced the reduction of neuronal NRG1 release involved in the pathogenesis of MHE.

We found that NRG1 is significantly downregulated, and that NRG1administration induced the restoration of the synaptic dysfunction seen in MHE brains. The addition of NRG induced a further increase in presynaptic marker (syntaxin) and post-synaptic marker (Homer) levels. FGF2 administration triggered the activation of NRG1/ErbB4 signaling in MHE rats. The reduction of FGF2 induced synaptic dysfunction and led to cognitive impairment via the inhibition of NRG1. We found that FGF2 may act through NRG1 activation, and regulation of NRG1 is important for synaptic spine formation. Furthermore, we found that NRG1 is a critical factor for inducing FGF2 secretion. Taken together, these findings indicate that exogenous decreased FGF2 may be associated with the reduction of NRG1 expression under MHE conditions, and both of FGF2 and NRG1 led to memory improvement.

Mammalian Sirt1, characterized as a NAD+-dependent histone deacetylase (*37*), is expressed in the hypothalamus (*38*). Moreover, mounting evidence implicates that Sirt1 is identified as a critical factor for normal synaptogenesis and cognitive function (*26, 39*). Therefore, Sirt1 is indispensable for cellular mechanisms underlying synaptic formation in rats. Our results demonstrate that FGF2 appears to facilitate the expressions of synaptic proteins via Sirt1, it is likely that FGF2 modulates learning and memory function through NRG1/ErbB4/Sirt1 signaling.

## Conclusions

our study demonstrated that FGF2 regulates dendritic spine generation and synaptic formation via NRG1 release and Sirt1 expression. Our findings also showed that FGF2 expression is decreased in MHE brain, and that FGF2 treatment mitigates the disruption of synaptic formation and memory impairment via neuronal NRG1/ErbB4 signaling. These findings highlight FGF2 as a promising potential treatment for MHE.

## List of abbreviation

(MHE): Minimal hepatic encephalopathy
(FGF2): Fibroblast growth factor-2
(NRG1): neuregulin 1
(CNS): central nervous system
(FGFR1): FGF receptor-1
(AD): Alzheimer’s disease
(WFT): water-finding task
(YM): Y-maze
(TAA): thioacetamid
(EL, the elapsed times for entry into the alcove): Entry latency,
(CL, the elapsed times for the first touching/sniffing/licking of the water tube): contacting latency
(DL, the elapsed times for the initiation of drinking from the water tube): drinking latency
(PHNs): Primary hippocampal rats neurons
(PCNs): Primary cortical rats neurons

## Availability of data and materials

The datasets used and/or analysed during the current study are available from the corresponding author on reasonable request.

## Authors’ contributions

Qichuan Zhuge and Saidan Ding contributed to the conception of the study.Jian Wang and Weishan Zhuge contributed significantly to analysis and manuscript preparation;Xiaoai Lu and Ruimin You performed the data analyses and wrote the manuscript;Leping Liu, He Yu, Yiru Ye, Xuebao Wang helped perform the analysis with constructive discussions. All authors read and approved the final manuscript.

## Acknowledgements

This study was funded by National Natural Science Foundation of China (81671042, 81300308, 81171088, 81371396).

---

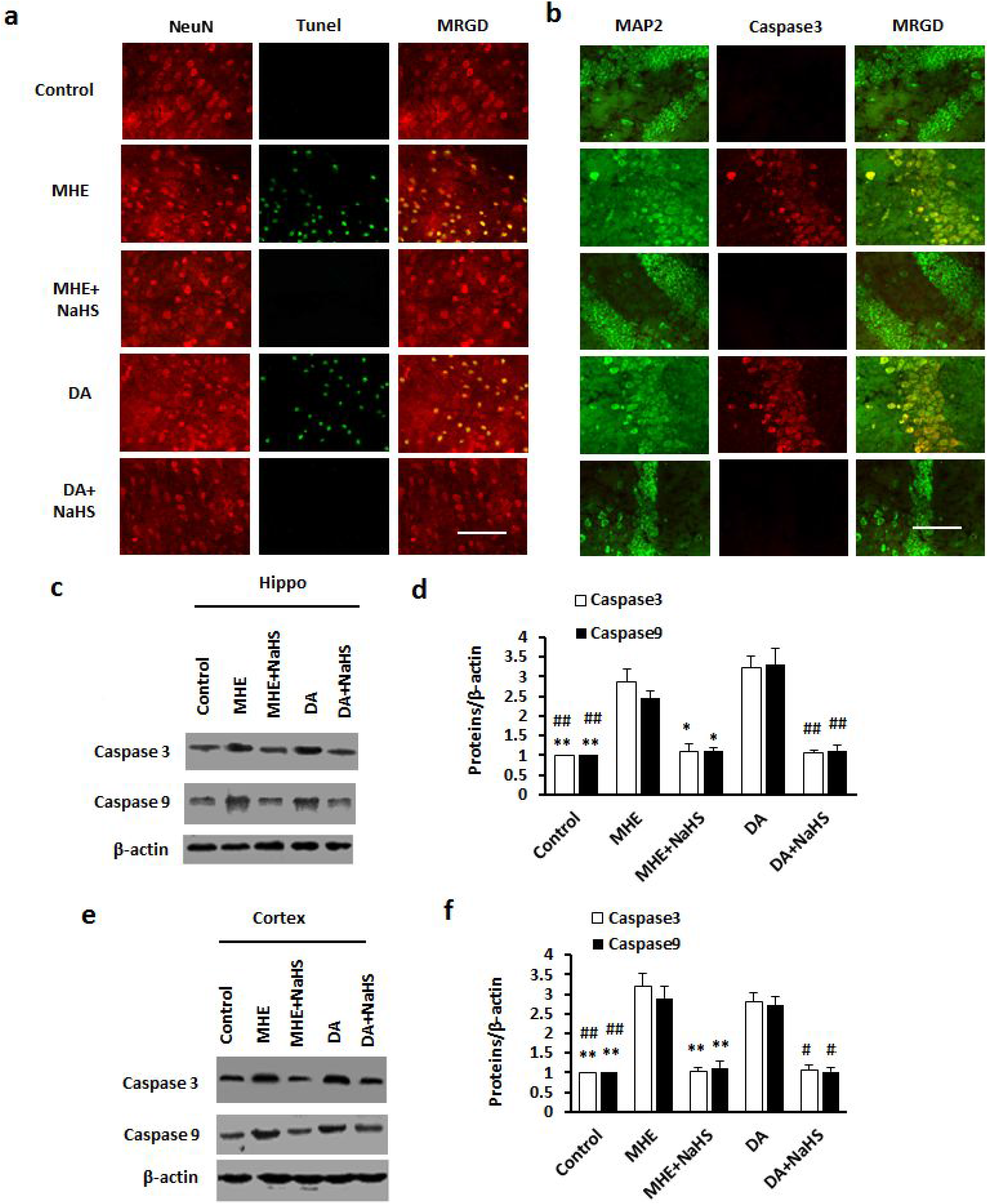

